# Mapping the integration of sensory information across fingers in human sensorimotor cortex

**DOI:** 10.1101/2021.07.07.451552

**Authors:** Spencer A. Arbuckle, J. Andrew Pruszynski, Jörn Diedrichsen

## Abstract

The integration of somatosensory signals across fingers is essential for dexterous object manipulation. Previous experiments suggest that this integration occurs in neural populations in the primary somatosensory cortex (S1). However, the integration process has not been fully characterized, as previous studies have mainly used two-finger stimulation paradigms. Here, we addressed this gap by stimulating all 31 single- and multi-finger combinations. We measured population-wide activity patterns evoked during finger stimulation in human S1 and primary motor cortex (M1) using 7T functional magnetic resonance imaging (fMRI) in female and male participants. Using multivariate fMRI analyses, we found clear evidence of unique non-linear interactions between fingers. In Brodmann area (BA) 3b, interactions predominantly occurred between pairs of neighbouring fingers. In BA 2, however, we found equally strong interactions between spatially distant fingers, as well as interactions between finger triplets and quadruplets. We additionally observed strong interactions in the hand area of M1. In both M1 and S1, these non-linear interactions did not reflect a general suppression of overall activity, suggesting instead that the interactions we observed reflect rich, non-linear integration of sensory inputs from the fingers. We suggest that this non-linear finger integration allows for a highly flexible mapping from finger sensory inputs to motor responses that facilitates dexterous object manipulation.

## Introduction

When writing with a pen or manipulating a Rubik’s cube in one hand, the sensorimotor system needs to integrate somatosensory information from multiple fingers to estimate the object’s shape, position, and movement within the hand. The mechanism that underlies this integration, however, remains poorly understood. We hypothesized that, in order to support flexible behavioural responses to any pattern of sensory stimulation across the hand, sensory inputs from neighbouring and non-neighbouring fingers need to be integrated in a non-linear fashion. This non-linear code provides the neural substrate necessary to detect any specific pattern of stimulation across the hand and forms the basis for learning flexible mappings between sensory inputs and motor responses of the hand.

Stimulation to the fingers is relayed from mechanoreceptors via the cuneate nucleus to the thalamus, with signals from different fingers remaining largely segregated (Florence, Wall, & Kaas, 1988, 1989). Signals from different fingers begin to interact substantially only in S1 and M1 (Hsieh et al., 1995). Cortical sensory processing is critical for dexterous hand control, as perturbing either the transmission of somatosensory information from the hand to the cortex (Moberg, 1958; Monzée, Lamarre, & Smith, 2003; Chemnitz, Dahlin, & Carlsson, 2013), or lesioning S1 (Carlson, 1981; Hikosaka et al., 1985; Brochier et al., 1999) grossly impairs fine manual dexterity. We refer here to S1 and M1 collectively as sensorimotor cortex. In the primate brain this comprises six cytoarchitectonically distinct Brodmann areas (BA): 4a, 4p, 3a, 3b, 1, and 2 (Brodmann, 1909; Powell & Mountcastle, 1959; Geyer et al., 1996). Inputs from the thalamic somatosensory nuclei vary across these regions, with BA 3a and BA 3b receiving most of the inputs, BA 4a and BA 4p receiving a substantial amount, and BA 1 and BA 2 receiving progressively fewer (Jones & Powell, 1970; Jones, 1975; Shanks & Powell, 1981; Darian-Smith & Darian-Smith, 1993). In nonhuman primates, neurons in BA 3b have receptive fields mainly devoted to single fingers, whereas in BA 1 and BA 2, receptive fields encompass multiple fingers (Hyvärinen & Poranen, 1978b; Sur, 1980; Iwamura et al., 1993). Measuring the coarse spatial organization for fingers in these regions with fMRI reveals comparable findings in humans, with finger representations becoming more spatially overlapping in posterior subregions of S1 (Krause et al., 2001; Martuzzi, et al., 2014; Besle et al., 2014). At the single-neuron level, paired finger stimulation generally results in lower firing rates relative to what would be expected from summing the firing rates resulting from single finger stimulation (Friedman, Chen, & Roe, 2008; Lipton et al., 2010; Reed et al., 2010; Thakur, Fitzgerald, & Hsiao, 2012). Together, these findings have been interpreted as evidence that inputs from multiple fingers are integrated in the sensorimotor cortex (Iwamura, 1998; Yau et al., 2016).

Everyday object manipulation often demands the integration of information across all fingers of the hand. In contrast, most previous studies have typically examined stimulation of only a few pairs of fingers. Consequently, the full nature of the interactions that occur between somatosensory inputs from all five fingers is not well characterized. Furthermore, it is unclear whether these previously reported suppressive interactions reflect the encoding of specific patterns of multi-finger stimulation (i.e., non-linear finger integration) or simply divisive normalization (Carandini & Heeger, 2011; Brouwer et al., 2015), where the inputs coming from individual fingers are linearly combined, but the net activity is suppressed through a diffuse inhibition. Studies of finger integration in humans also share these limitations (Gandevia, Burke, & McKeon, 1983; Hsieh et al., 1995; Biermann et al., 1998; Ishibashi et al., 2000; Hoechstetter et al., 2001; Ruben et al., 2006).

Here, we addressed this gap by studying all 31 possible finger combinations by simultaneously stimulating one, two, three, four, or five fingers of the right hand. We measured activity patterns evoked in the hand area of the sensorimotor cortex using 7T fMRI while human participants experienced passive tactile stimulation. Consistent with previous work, we found progressive overlap of single finger representation in sensorimotor cortex. By analyzing the multivoxel activity patterns in each subregion, we also found clear evidence for progressively stronger multi-finger interactions in posterior S1 and M1.

## Materials and Methods

### Participants

Ten healthy participants were recruited for the study (7 males and 3 females, mean age=25.5, SD=3.24; median Edinburgh handedness score=80.0, median absolute deviation=20.0). Participants completed one training session and two experimental sessions. During the training session, participants were familiarized with the finger stimulation task. In the two experimental sessions, participants experienced finger stimulation while undergoing 7T fMRI. All participants provided informed consent before the beginning of the study, and all procedures were approved by the Office for Research and Ethics at the University of Western Ontario.

### Apparatus

We used a custom-built five-finger keyboard to apply stimulation independently to each of the five fingers of the right hand. Each finger was comfortably restrained above an immobile key using a clamp covered with padding (Fig. 1A). The clamp prevented any hand or finger movement and ensured that the passive stimulation was delivered to a constant area of the fingertip. We delivered independent stimulation to each fingertip using a pneumatic air piston mounted underneath each key. The pistons were driven by compressed air (100 psi) delivered from outside the MRI scanning room through polyvinyl tubes. The forces applied to the fingertips were monitored using force transducers (Honeywell-FS series, dynamic range=0-16N, resolution<0.02N, sampling rate=200Hz), and the air pressure for each piston rod was independently regulated using PID control to deliver forces of ~3 Newtons to each fingertip (one participant experienced stimulation of ~2N). The piston rods (diameter=3mm) deformed the skin of the fingertip. As the padding prevented movement of the finger, the stimulation was predominantly tactile in that it involved deformation of the skin.

**Figure 1.**
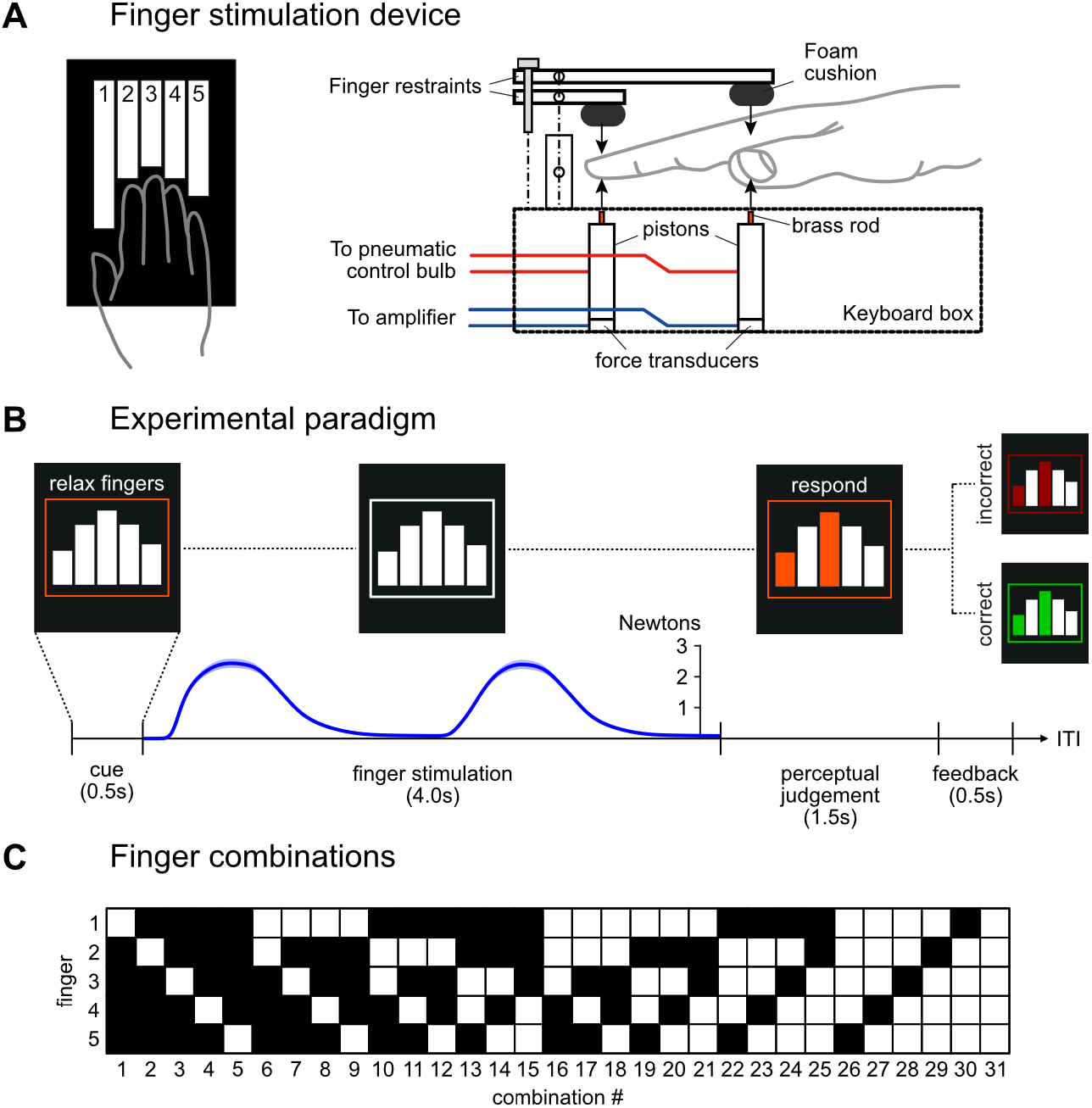
Experiment design. (**a**) Participants experienced tactile stimulation using a custom-built finger stimulation box. Each finger was independently restrained, and pneumatic pistons were used to deliver stimulation to each fingertip. (**b**) Schematic illustration of a single trial (ITI: inter-trial-interval). The blue trace shows the mean finger stimulation force (±SEM across participants), time aligned to the start of the stimulation phase. (**c**) Participants experienced stimulation of all 31 possible single- and multi-finger combinations.

### Task

#### Finger stimulation

While lying in the scanner, participants viewed a back-projection screen through a mirror mounted to the head coil. They saw five bars in the centre of the screen, surrounded by a box (Fig. 1B). Each bar corresponded to one of the five fingers of the right hand (left-to-right: thumb-to-little finger). All 31 possible finger combinations were stimulated (5 single-finger, 10 two-finger, 10 three-finger, 5 four-finger, and 1 five-finger configuration; Fig. 1C).

Each trial lasted 6.5s and consisted of four phases (Fig. 1B): a cue phase (0.5s), finger stimulation (4s), perceptual judgement (≤1.5s), and feedback (≥0.5s). The cue phase alerted participants to the start of the trial. During the cue, the outline box turned orange and the words “RELAX FINGERS” where presented on screen, instructing participants to relax their hand. No information was provided about which finger combination was going to be stimulated, and therefore participants remained naïve about the stimulation until it occurred. At the start of the stimulation phase, the words disappeared from the screen and the box turned white, after which one of the 31 possible finger combinations was stimulated. The stimulated force on each finger approximated a rectified sine wave, gradually increasing and decreasing. Each “wave” of stimulation lasted approximately 1s, and two complete waves were delivered during each trial. Across all fingers and combinations, the average measured peak force was 2.67±0.17N.

#### Mismatch detection

In the perceptual judgement phase of each trial, we presented a visual arrangement of a specific finger combination, with the boxes corresponding to the stimulated fingers turning orange (Fig. 1B). To ensure that participants remained attentive during the experiment, we asked them to detect the occurrence of relatively rare (5% of trials) mismatches between the visually presented and stimulated patterns (probe trials). Participants were asked to detect this mismatch and indicate it by pressing their right thumb (2N threshold). If the presented and stimulated pattern matched, they were instructed to refrain from pressing any finger. We chose this particular task because it ensured that participants remained attentive, but it did not explicitly require the integration of sensory information across fingers. On the contrary, the task demanded that sensory information from each finger be analyzed separately.

Participants had 1.5s to judge and respond. After either 1.5s elapsed (indicating a match) or immediately following a thumb press (indicating a mismatch), participants were provided feedback on their response by changing the colour of the finger combination green (correct) or red (incorrect). The feedback was presented on-screen for ≥0.5s, such that the feedback and response phases together lasted 2s regardless of response type.

To encourage good performance, participants received points based on the performance of their perceptual judgements. They were awarded 1 point for correctly identifying a matching configuration, and 10 points for correctly identifying a mismatched configuration. False alarms were penalized by −1 point, and misses (failing to recognize a mismatch configuration) were penalized by −10 points. Points were combined across imaging runs and used to calculate a financial bonus. Behavioural performance on the perceptual judgement task was high (overall error rate=1.77±0.40%) with participants being well able to discriminate perceptual mismatch trials (hit rate=86.97±5.09%, false alarm rate=1.20±0.21%, d’=3.68±0.31). Although participants tended to be conservative in their response behaviour (i.e., they were somewhat biased to not report a mismatch: c=0.44±0.11), this was expected because most of the trials (95%) were matches and thus required participants to refrain from making a thumb press. Together, this resulted in participants making a thumb press on 5.35±0.20% of all trials.

### Procedure

Participants completed one training session and two imaging sessions. Each session was comprised of several runs. Each run contained 62 trials, with two trials for each of the 31 finger combinations (see Finger stimulation). 5% of these trials contained perceptual mismatches (see Mismatch detection). Trials were separated by a variable inter-trial-interval (ITI), drawn randomly from the set of possible ITIs {1s, 2.5s, 4s, 5.5s, 7s, 8.5s, 10s}, with the probability p=[0.37, 0.24, 0.16, 0.10, 0.06, 0.04, 0.03] for each ITI, respectively. Thus, shorter delays occurred more often and longer delays occurred less often. The order of trials, including the position of the mismatch trials, was randomized across runs and participants.

During training, participants performed runs of trials until they achieved an overall error rate of 0% in one run. Participants completed the training session 1 −2 days before the first scanning session. For the imaging sessions, participants completed 11 total runs, yielding 682 total trials (31 combinations × 2 repeats × 11 runs). These 11 runs were split over two separate scanning sessions for each participant, usually within the same week.

### MRI data acquisition

We used high-field functional magnetic resonance imaging (fMRI, Siemens 7T Magnetom with a 32-channel head coil at Western University, London, Ontario, Canada) to measure the blood-oxygen-level dependent (BOLD) responses evoked in participants. Each participant completed 11 runs of trials across two separate scanning days, usually with 6 runs on the first day and 5 runs on the second day. Each run lasted 614s. During each run, 410 functional images were obtained using a multiband 2D-echoplanar imaging sequence (GRAPPA, in-plane acceleration factor=2, multi-band factor=3, repetition time [TR]=1500ms, echo time [TE]=20ms, in-plane resolution 148 × 148 voxels). Per image, we acquired 66 interleaved slices (without gap) with isotropic voxel size of 1.4mm. The first 2 images in the sequence were discarded to allow magnetization to reach equilibrium. To estimate magnetic field inhomogeneities, we acquired a gradient echo field map at the end of the scanning session on each day. Finally, a T1-weighted anatomical scan was obtained using a magnetization-prepared rapid gradient echo sequence (MPRAGE) with a voxel size of 0.75mm isotropic (3D gradient echo sequence, TR=6000ms, 208 volumes).

### fMRI preprocessing and first-level analysis

Functional images were first realigned to correct for head motion during the scanning sessions (3 translations: x,y,z; 3 rotations: pitch, roll, yaw), aligned across sessions to the first image of the first session, and co-registered to each participant’s anatomical T1-image. Within this process, we used B0 fieldmaps from each imaging session to correct for image distortions arising from magnetic field inhomogeneities (Hutton et al., 2002). Due to the relatively short TR (1.5s), no slice-timing correction was applied, and no spatial smoothing or normalization to a standard template was applied.

The minimally preprocessed data were then analyzed using a general linear model (GLM; Friston, Jezzard, & Turner, 1994) using SPM12 (fil.ion.ucl.ac.uk/spm/) with a separate regressor for each of the 31 possible finger combinations in each run. The activation during stimulation was modeled using a boxcar function that spanned the stimulation phase of each trial, convolved with a hemodynamic response function with a delayed onset of 1.5s and a poststimulus undershoot at 12s. Given the low error rate, all trials were included in the analysis, regardless of the perceptual judgement accuracy. To capture activity evoked by the thumb press during mismatch detection, we included one thumb press regressor in each run which modeled all thumb responses per run (duration: 1s). Because the response was always the same thumb press and occurred with roughly equal probability for each finger combination, any incomplete modelling of the response-evoked activity should not influence our results pertaining to the differences in activity patterns between finger combinations. Finally, we included an intercept regressor for each run, yielding 363 total regressors (33 regressors × 11 runs).

To model the long-range temporal autocorrelations in the functional timeseries, we used the SPM FAST autocorrelation model. High-pass filtering was then achieved by temporally prewhitening the functional data with this temporal autocorrelation estimate. This analysis yielded one beta-weight for each voxel for each of the 31 finger combinations per run for each participant. Collectively, these defined the estimated activity patterns. We did not further analyze the activity pattern from the thumb press regressor. From these beta-weights, we calculated the average percent signal change (PSC) evoked by each finger combination relative to the baseline for each voxel, yielding 31 PSC values per voxel.

### Surface-based analyses

#### Surface reconstruction

We used Freesurfer (Fischl, Sereno, & Dale, 1999) to reconstruct the cortical surface from the anatomical image of each participant, with each node having a location on the white-matter/gray-matter surface and the pial surface. The surfaces were then spherically registered to match a template atlas (FreeSurfer’s Left-Right 164k node template) based on a sulcal-depth map and local curvature.

#### Projection of activity patterns to cortical surface

To visualize the evoked activity patterns, the individual patterns were projected on the individual surface. For each surface node, all voxels that lay between the white matter and the pial surface location for that node were averaged. To avoid the mixing of signals between M1 and S1 across the central sulcus, we excluded voxels that projected to two dis-connected groups of surface nodes (one on the anterior and one on the posterior bank of the central sulcus) with the second projection accounting for at least 25% of the total surface (see github.com/DiedrichsenLab/surfAnalysis for details).

#### Surface-based searchlight

For multivariate analysis, we used a surface-based searchlight approach (Oosterhof et al., 2011). For each node on the individual surface reconstruction, we used a geodesic distance metric to define a circular region on the surface that included the nearest 160 voxels between the pial and white/gray-matter surface (average geodesic radius=5.85±0.04mm). The use of a geodesic metric ensured that each searchlight region did not include voxels across a sulcus. The set of activity patterns from these voxels were then analyzed together (see Multivariate fMRI analysis) and the results were assigned to the corresponding centre node. The searchlight analysis was primarily used for visualization purposes (e.g., Fig. 2C).

**Figure 2.**
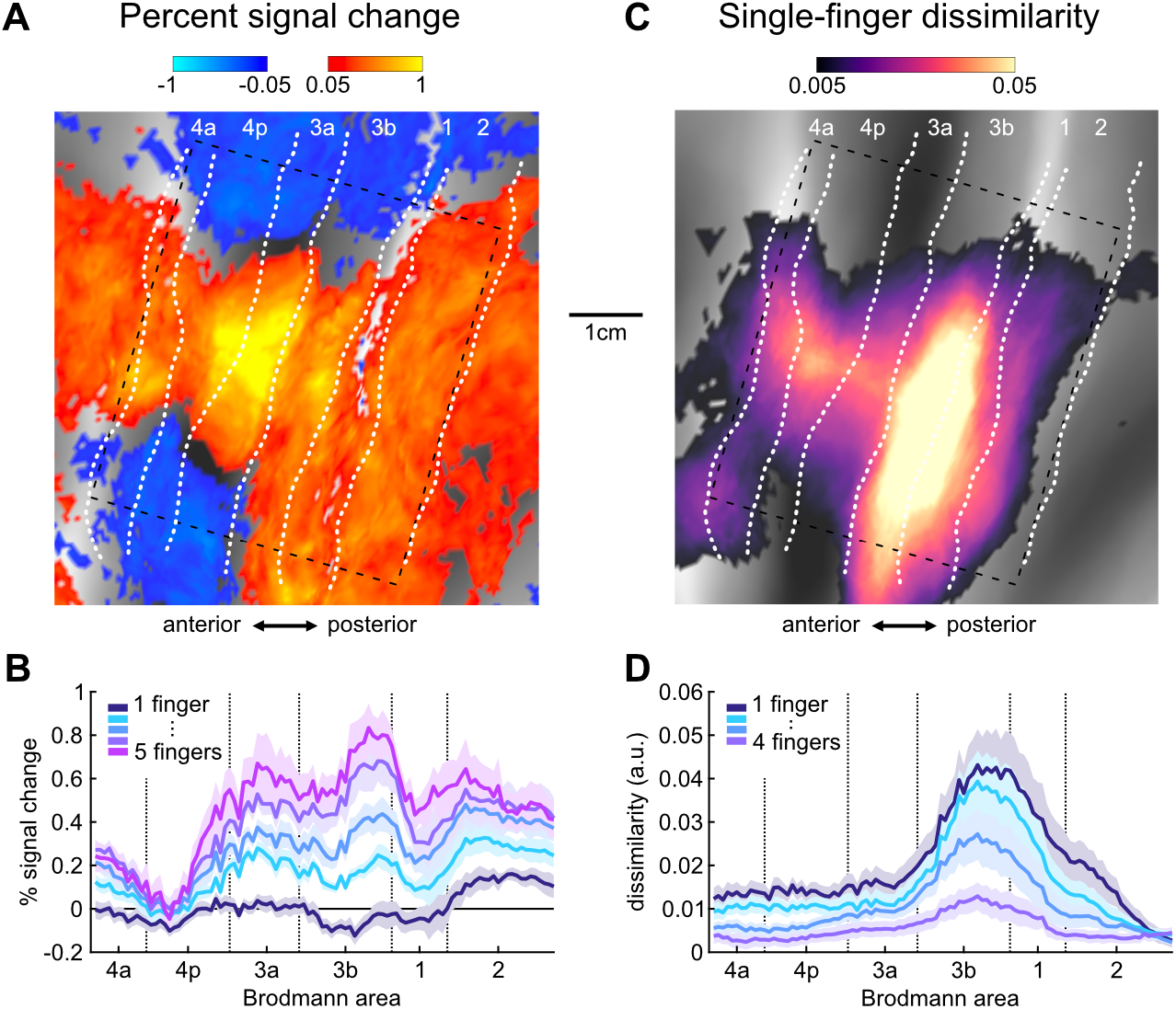
Activation and representation of fingers in sensorimotor cortex. (**a**) Group-average percent signal change (relative to resting baseline) across all 31 possible finger combinations, projected to a flattened view of the left cortical sensorimotor areas around the central sulcus. Approximate boundaries of cyctoarchitectonic areas (Fischl et al., 2008) are indicated by white dotted lines. The gray background indicates the average sulcal depth, with gyri in light, and sulci in dark colors. The rectangle indicates the area of averaging in the cross-sections in B and D. The scale bar approximates 1cm on the flattened surface. (**b**) Cross-sectional profiles (see Methods) of the average percent signal change (±SEM across participants) within the dashed rectangle in A, grouped by the number of fingers in each combination. The x-axis corresponds to the approximate spatial location along the rostral-caudal axis spanned by the rectangular box on the cortical surface. Vertical dashed lines mark the approximate boundaries between Brodmann areas. (**c**) Cortical surface map of the average crossnobis dissimilarity between activity patterns evoked by single-finger stimulation. (**d**) Cross-sectional profiles of the average crossnobis dissimlarity (±SEM across participants) between pairs of single-finger, 2-finger, 3-finger, or 4-finger combinations.

#### Cross-sectional profile plots

To create the cross-sectional surface profiles of univariate and multivariate measures (e.g., Fig. 2B & D), we used the data that fell into a rectangular box that spanned the sensorimotor cortex of each participant (dashed rectangle in Fig. 2A). We orthogonally projected surface nodes within the rectangular box onto a cross-sectional line that approximately spanned the rostral-caudal axis from BA 4a to 2. Following, data from the projected nodes where then averaged within 101 equidistant bins along this cross-section. Figure 2B and D show the group-averaged cross-sectional profiles. As with the searchlight analysis, the cross-sectional profiles were primarily for visualization purposes.

#### Surface-based tessellation

To conduct more computationally intensive representational model comparisons (see Representational model analysis) across the cortical surface, we employed a coarser alternative to the continuous surface-based searchlight approach: We defined a reduced set of surface patches by tessellating the left hemisphere into 1442 regular hexagonal tessels. We then sub-selected a set of tessels that had enough reliable differences between stimulation conditions to allow for model-comparison. Specifically, a tessel was included if the group-averaged continuous searchlight result showed an average pattern dissimilarity across all activity patterns of ≥0.005. This criterion yielded 82 tessels that spanned the surface of sensorimotor cortex, with an average of 98.93±2.81 voxels per tessel.

### Regions of interest definition

For a targeted analysis of subregions of the sensorimotor cortex, we used a probabilistic cytoarchitectonic atlas projected to the cortical surface (Fischl et al., 2008) to define Brodmann areas 4a, 4p, 3a, 3b, 1, and 2. Surface nodes were assigned to the region that had the highest probability, and this probability needed to exceed 0.25. We further restricted these regions to the hand area by excluding nodes that fell 2.5cm above and 2.5cm below the hand knob anatomical landmark (Yousry et al., 1997). To avoid smearing activity across the central sulcus, we excluded (as in the surface projection) voxels with >25% of their volume in the gray matter on the opposite side of the central sulcus. This yielded 546.70±35.45 voxels for BA 4a, 642.20±41.99 voxels for BA 4p, 275.20±10.48 voxels for BA 3a, 711.00±31.05 voxels for BA 3b, 602.40±33.50 voxels for BA 1, and 1034.30±57.80 voxels for BA 2.

### Single-finger selectivity

#### Voxel selection

To quantify the selectivity of each voxel for a specific finger, we considered only the activity estimates for the single-finger conditions. We selected voxels from each region that showed significant modulation (relative to baseline) during any single-finger stimulation, by conducting an omnibus *F*-test per voxel, retaining only voxels that were significant on an p=0.05 level (uncorrected). This criterion selected 8.98±1.24% of the voxels from BA 4a (50.70±8.58 voxels), 7.62±0.88% from BA 4p (48.30±5.60 voxels), 8.19±1.23% from BA 3a (22.50±3.61 voxels), 14.84±2.10% from BA 3b (105.60±15.73 voxels), 16.60±2.57% from BA 1 (101.50±16.78 voxels), and 10.91±1.46% from BA 2 (116.50±20.91 voxels). We verified in simulations that this voxel selection approach did not bias the subsequent selectivity analysis, but simply increased its power. This is because the omnibus *F*-test only tests *if a* voxel is tuned to one or more fingers, whereas the selectivity analysis characterizes the *shape* of the voxel’s tuning to different fingers.

#### Quantifying selectivity

We then normalized the tuning curves (5 activity values for each voxel), such that the maximal response equaled 1 and the lowest response equaled 0. For the plots in Figure 3B, we grouped the voxels according to the most preferred finger.

**Figure 3.**
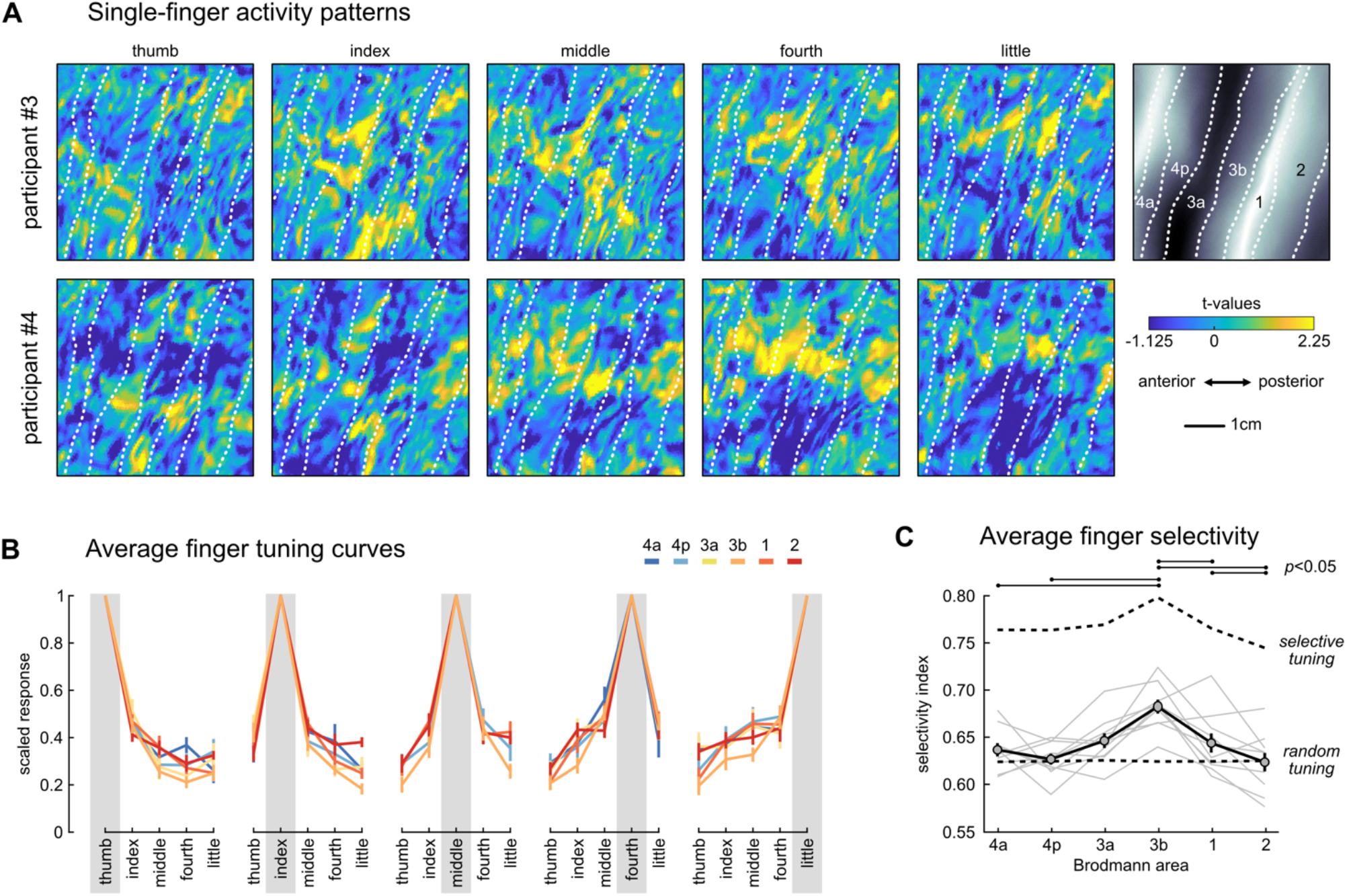
Finger tuning in sensorimotor cortex. (**a**) Activity patterns for each of the five fingers from one participant, projected onto a flattened cortical surface and cut to include BA 3a to BA 2 in each panel. (**b**) Average scaled voxel tuning curves arranged by most preferred finger (denoted by the gray box). Each colour corresponds to different regions. (**c**) Finger selectivity coefficients per region. Light gray lines reflect selectivity coefficients per participant, and solid black line reflects the average across participants. The lower dashed line reflects the average expected selectivity if voxels were randomly tuned to fingers, while the upper dashed line reflects the average expected selectivity if voxels only responded to a single finger. Expected values take into account the empirical noise variance in each region and participant. A-priori paired t-tests were conducted between normalized selectivity coefficients (see Methods) from different regions, and lines above the plot denote significant differences. Error bars in B and C reflect SEM across participants in each region.

Using the normalized tuning curve for each voxel, we calculated the voxel’s selectivity (*λ*) as:

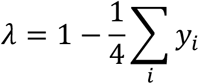

where *y_i_* are the normalized responses to the four “less-preferred” fingers. This yields the average difference between the activity evoked by the finger that the voxel is most *tuned* to (the maximal activity) and all other finger activities. Therefore, *λ* = 1 indicates a voxel is only active during stimulation of a specific finger. Conversely, *λ* < 1 indicates that a voxel also responds to stimulation of other fingers. For each participant, we averaged the resulting selectivity coefficients across the selected voxels per region. This yielded one selectivity coefficient per region per participant, which are plotted in Figure 3C.

#### Controlling for measurement noise

Due to measurement noise, the estimated selectivity coefficients will always be <1, even if all voxels would only respond to a single finger. The level of signal-to-noise may differ across regions and participants, making it inappropriate to directly compare the raw selectivity coefficients. Furthermore, even completely random tuning would still result in estimated selectivity coefficients >0. To address this issue, we simulated voxel tuning curves under two different generative models. First, for random tuning, we simulated voxels with tuning that was drawn from a multivariate Gaussian distribution, with covariance identical to the group-averaged finger-by-finger correlation matrix (Ejaz, Hamada, & Diedrichsen, 2015). Second, for highly selective tuning, we simulated voxels that were selective for stimulation of a single finger and remained unresponsive to all other fingers. Both simulations were scaled, so that the average diagonal of the covariance matrix matched the signal strength for that region and participant 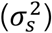. We then added the measurement noise, drawn from a normal distribution with variance set to 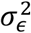, again matched to that region and participant.

To estimate 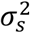 and 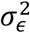 from the real data of each participant, we first vectorized the matrix of mean-centred tuning curves for each run, and then calculated the average covariance between these vectors across runs. An estimate of 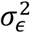 could then be obtained via 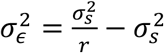, where *r* is the average Pearson’s correlation between the vectorized tuning curves across runs. This expression arises because we assume the noise in the real data is independent (i.e., uncorrelated) across different runs, and therefore, the expected value of the Pearson’s correlation between vectorized tuning curves from different runs is 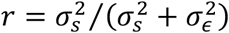.

Using these estimates, we simulated 1000 datasets with random tuning and 1000 datasets with perfectly selective tuning for each region in each participant, each with the same number of voxels as the real data. All data sets were simulated with 11 imaging runs. As for the real data, we then applied the voxel selection to the simulated data. We calculated the average selectivity across voxels in each simulated dataset and averaged the selectivity coefficients across simulations. For statistical comparison of selectivity coefficients across regions, the selectivity of the real data was then normalized such that a selectivity of 0 reflected the expected value under random tuning and 1 reflected the expected value for highly selective tuning. This normalization was done for each region and participant separately and ensured that comparisons across regions were not biased by differences in noise.

### Representational similarity analysis

To test for reliable differences between fMRI activity patterns for each condition (i.e., the first-level GLM beta-weights), we used the cross-validated squared Mahalanobis dissimilarity (i.e., crossnobis dissimilarity, Walther et al., 2016). Cross-validation ensures the dissimilarity estimates are unbiased, such that if two patterns differ only by measurement noise, the mean of the estimated dissimilarities would be zero. This also means that estimates can sometimes become negative (Diedrichsen et al., 2020; Diedrichsen, Provost, & Zareamoghaddam, 2016). Therefore, dissimilarities significantly larger than zero indicate that two patterns are reliably distinct, and we take this as evidence that features about the finger combinations are represented in the activity patterns.

The crossnobis dissimilarity *d* between the fMRI activity patterns (***x***) for conditions *i* and *j* was calculated as:

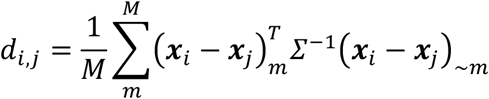

where the activity patterns from run *m* are multiplied with the activity patterns averaged over all runs except *m* (~*m*). *Σ* is the voxel-wise noise covariance matrix, estimated from the residuals of the first-level GLM, and slightly regularized to ensure invertibility. Multivariate noise-normalization removes spatially correlated noise and yields generally more reliable dissimilarity estimates (Walther et al., 2016). Analyses were conducted using functions from the RSA (Nili et al., 2014) MATLAB toolbox. For the searchlight results (Fig. 2C & 2D), we averaged the resulting dissimilarities based on whether they were between single-finger patterns, 2-finger patterns, 3-finger patterns, or 4-finger patterns.

### Representational model analysis

We used representational models to infer what feature sets were present in the activity patterns from each region. A representational model characterizes the tuning curves of a group of voxels or neurons. In the sense we are using it here, it specifies a probability distribution over all possible tuning curves (Diedrichsen & Kriegeskorte, 2017). Here, we used an encoding-style approach (Naselaris et al., 2011) to specify and evaluate representational models that predicted activity patterns for all finger combinations using various feature sets. Models were fit and evaluated using the PCM toolbox (Diedrichsen, Yokoi, & Arbuckle, 2018), using crossvalidation across imaging runs for each region in every participant.

#### Linear Model

The linear model predicted that activity patterns evoked during multi-finger stimulation were the sum of the constituent single-finger patterns:

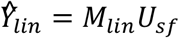

where 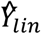 are the [31 × P voxels] predicted activity patterns, *U_sf_* is a [5 features × *P* voxels] matrix of single-finger feature patterns, and *M_lin_* is a [31 combinations × 5 features] indicator matrix denoting which finger(s) are in each of the combinations. To complete the representational model, we also specified that the single-finger features had a second-moment matrix (co-variance matrix without subtraction of the mean across voxels) of *G_lin_*. The second moment matrix of finger-related patterns is highly invariant across individuals, reflecting the high correlations of patterns from neighbouring fingers, and the low correlation of the pattern of the thumb with the other fingers (Ejaz et al., 2015; Arbuckle et al., 2020). We determined the exact form of the matrix for each region separately, using the group-averaged second moment matrix 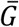 from the region under analysis. Specifically, we determined the second moment matrix for the single-finger patterns that would best approximate the overall second moment matrix 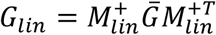, where 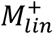 is the Moore-Penrose pseudoinverse.

#### Multi-finger interaction models

We also constructed 3 multi-finger interaction models. Like the linear model, these models assumed that the multi-finger patterns were the sum of the constituent single finger patterns, but also included specific interaction effects between specific fingers. These models took the same general form as the linear model above. For the 2-finger interaction model, we included 10 terms indicating the presence of a specific pair of fingers (i.e. when the pair of fingers was pressed, the regressor was 1 and 0 otherwise), in addition to the 5 model features for the individual fingers. In the 3-finger interaction model, we additionally added 10 regressors indicating the presence of each unique 3-finger combination. Finally, the 4-finger model included, in addition to the 3-, 2-, and 1 -finger terms, the five possible 4-finger interactions, resulting in 30 model features overall. For each of the models, the second-moment matrix for the feature patterns *U* was estimated from the group-averaged second-moment matrix as for the linear model.

#### Adjacent and non-adjacent 2-finger interaction models

To test the strength of finger interactions between adjacent and non-adjacent finger pairs, we created two modified versions of the 2-finger interaction model. In the first version, we only included the 4 adjacent finger-pairs as interaction terms. In the second version we included only the 6 non-adjacent finger pairs.

#### Linear-nonlinear model

The linear-nonlinear model predicted that activity patterns for single-fingers combined linearly when multiple fingers were stimulated, but that the overall activity was compressed by a unknown, non-linear function *f*, 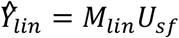. Such a non-linearity could arise from a global divisive normalization of neural activity in the region, or from non-linearities in the relationship between neural activity and the BOLD signal. To approximate any form of such compressive non-linearity, we created a model, based on the linear model, that allowed for flexible scaling of the predicted multi-finger patterns. All predicted patterns that included the same number of stimulated fingers were scaled by a common factor. These scaling factors, as well as the single-finger feature patterns *U_sf_*, were estimated from the training data.

#### Null-model

As a baseline for model comparison, we defined a null-model that predicted overall activity scaled with the number of fingers stimulated, but that the patterns lacked finger-specificity. Under this model, the predicted patterns were derived from the average activity patterns for the single-finger, 2-finger, 3-finger, 4-finger, and 5-finger combinations, respectively. For example, the predicted single-finger patterns was the average pattern across the five single-finger conditions from the training data.

#### Noise-ceiling model

To provide an estimate of how much systematic variation could be explained in the data given measurement noise, we included a “noise-ceiling” model. The predictions under the noise-ceiling model were simply the 31 activity patterns from the training data. Note that this fully saturated model differs from the 4-finger interaction model only by the addition of a single model term that models the specific non-linearities arising during the stimulation of all 5 fingers. The second-moment matrix for this model was set to the observed group-averaged estimate 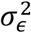 for the region under analysis.

#### Model fitting

We fit and evaluated the different models within each participant, using a leave-one-run-out cross-validation procedure. For each cross-validation fold, the training data were the activity patterns from all imaging runs except one, and the test data were the activity patterns from the left-out run.

For a representational model with the assumption that both noise and signal have multivariate Gaussian distribution (Diedrichsen & Kriegeskorte, 2017), the feature patterns for each model can be estimated from the training data *Y_train_* as:

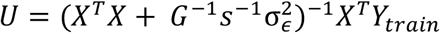

where *X* is a model-specific design matrix that denoted which feature(s) were present in each of the rows (activity patterns) in *Y_train_, G* is the model-dependent second-moment matrix as specified above, *s* indicates the strength of the signal in *Y_train_*, and 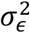 is the variance of the random run-by-run noise. Note that this estimation takes the form of linear regression with Tikhonov regularization. The strength of the regularization to the model prior was determined by 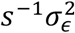. For each cross-validation fold, we derived the maximum-likelihood estimate of *s* and 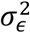 using the PCM toolbox. For the linear-nonlinear model, we additionally fitted the scaling parameters by minimizing the residual sums-of-squares of the model predicted patterns to the training data.

#### Model evaluation

The estimated feature patterns (*U*) were then used to predict activity patterns under the corresponding model. We compared the predicted 31 condition patterns with the left-out test data. The model fits were evaluated using Pearson’s correlation. For this, the predicted and test patterns were first mean-centred (per voxel), then correlated across all voxels and conditions within each cross-validation fold. We averaged the correlations across cross-validation folds to yield a single estimate of model performance per participant per region. We preferred Pearson’s correlation as our evaluation metric over the coefficient of determination, as it is less dependent on the exact choice of regularization coefficient (Diedrichsen & Kriegeskorte, 2017), and therefore provides a more robust evaluation.

Finally, the model fits were normalized between 0 and 1, using the fits of the null and noise-ceiling models. This normalization approach was necessary because, as illustrated by the fits of the noise ceiling models, measurement noise varied across regions (raw Pearson’s r in BA 4a=0.054±0.0063, BA 4p=0.054±0.0058, BA 3a=0.074±0.010, BA 3b=0.116±0.013, BA 1=0.074±0.011, BA 2=0.038±0.005). Normalization of the model fits enabled us to meaningfully compare fits across regions with varying levels of measurement noise. Normalized fits >0 indicated that the model captured more information than the simple scaling of overall activity (null model), and fits <1 indicated that there was structured variance in the activity patterns that remained unaccounted for in the model(s). The normalization of model fits was done for each region and participant separately.

### Experimental design and statistical analyses

All statistical tests were performed in MATLAB R2019a (Mathworks, Inc.). We set the significance level in our analyses to *p*=0.05. When a test involved multiple comparisons without any specified *a priori* hypotheses, we adjusted the significance level by dividing by the number of comparisons (i.e., Bonferroni correction). For clarity, we report uncorrected *p*-values in the text. The Bonferroni-corrected statistical threshold is reported as an *α*-value when appropriate. In cases where we had *a priori* hypotheses, we did not correct for multiple comparisons (i.e., replicating single-finger pattern overlap; Fig. 3C). To compare evoked activity, dissimilarities, or normalized model fits across regions, we used within-participant repeated measures ANOVAs. We used two-sided paired t-tests to compare the model fits to the fit of the noise-ceiling in each region. If the model fit did not differ significantly from the fit of the noise-ceiling model, we considered the model to be as good as the noise-ceiling. Therefore, to remain conservative, we evaluated uncorrected *p*-values and did not correct for multiple comparisons for this analysis, as this correction would lower the bar for what would be considered a “good-fitting” model.

### Data and Code Accessibility

The analyses reported in this paper relied on code from the Representational Similarity Analysis (github.com/rsagroup/rsatoolbox_matlab) and Pattern Component Modeling (github.com/jdiedrichsen/pcm_toolbox) MATLAB toolboxes. The pre-processed data and code necessary to reproduce analyses and plots are available on github (github.com/saarbuckle/finger-sensory-integration).

## Results

### Finger stimulation evokes broadly distributed activity in sensorimotor cortex

Using high-resolution 7T fMRI, we measured the activity patterns evoked by passive finger stimulation in the brains of 10 human participants. Stimulation was delivered independently to each fingertip of the right hand by indenting the skin with a small rod pushed by a pneumatic piston. We tested the entire set of 31 single- and multi-finger combinations. To keep participants engaged during the experiment, they were instructed to detect rare mismatches between the stimulated combination and a visual probe presented after finger stimulation (see Methods).

Figure 2A shows the group-average percent signal change (relative to rest) during righthand finger stimulation on a flattened surface view of the cortical hand regions in S1 and M1 of the left hemisphere. During single-finger stimulation, evoked activity was low, but as more fingers were stimulated, we observed an increase in overall activity across subregions of the sensorimotor cortex. To statistically evaluate activity, we subdivided the hand region into six anatomically defined Brodmann areas using a cytoarchitectonic atlas (Fischl et al., 2008), spanning from BA 4a to BA 2 (see Methods). In each BA subregion, activity increased when more fingers were stimulated (all *F_(4,36)_*≥4.730, all *p*≤0.0036; see Fig. 2B).

This activity increase does not provide a detailed view of how sensory information from different fingers is integrated in the human sensorimotor cortex. As a starting point to address this question, we quantified how dissimilar the local single-finger activity patterns were from each other. We used a crossvalidated estimate of the dissimilarity measure (crossnobis, see Methods), such that a value of zero indicated that patterns only differed by noise, and dissimilarity values greater than zero indicated that the patterns were distinct. The average dissimilarities showed that single-finger stimulation evoked distinct finger patterns in all subregions (Fig. 2C). Indeed, all considered BA regions showed highly significant finger-specific pattern differences (one-sided t-test>0: all *t_9_*≥3.012, all *p*≤0.0073, Bonferroni corrected *α*-value=0.0083), which suggests that each region received information about the stimulated fingers. Dissimilarities between all 2-, 3-, and 4-finger combinations showed a similar spatial distribution, although the overall magnitude of the dissimilarities was reduced compared to the single-finger patterns (Fig. 2D). This finding is expected because multi-finger combinations also share an increasing number of fingers.

### Increasing overlap of single-finger patterns in sensorimotor cortex

Based on previous electrophysiological (Hyvärinen & Poranen, 1978b; Sur, 1980; Iwamura et al., 1993) and fMRI (Martuzzi et al., 2014; Besle et al., 2014) results, we would expect to find relatively focal single-finger activation in BA 3b, with more overlap between fingers in other parts of the sensorimotor cortex. This seemed to be the case as shown in the single-finger patterns for two exemplary participants (Fig. 3A). Each finger activated a quite distinct region of BA 3b and BA 3a. In contrast, the spatial patterns for each finger in BA 1 and BA 2, as well as in M1 (BA 4a and BA 4p) were more complex and involved multiple “hot spots” per finger, with substantial overlap between fingers.

We quantified this observation by computing a measure of finger-selectivity for each voxel. We selected voxels from each region that were significantly responsive to stimulation of any individual finger (see Methods), and scaled the responses of these voxels, such that the activity associated with the finger that evoked the largest positive response (i.e., the most-preferred finger) equaled 1, and the activity associated with the finger that evoked the smallest response (i.e., the least-preferred finger) equaled 0. If the voxel was only tuned to one finger, all non-preferred fingers would have a value of zero. The average scaled responses for the 4 non-preferred fingers therefore can be used as a measure of the selectivity of that voxel (Fig. 3B). To then define a selectivity index, we subtracted the averaged scaled responses for the 4 non-preferred fingers from 1, such that 1 indicates maximal selectivity, and 0 equal activation for all fingers. We averaged the selectivity coefficients across voxels per participant in each region.

Before comparing the selectivity coefficients across regions, we needed to address one last problem – namely, regions with less reliable data could appear to be more broadly tuned to multiple fingers simply because higher measurement noise makes the tuning less clear. This is a concern because the strength of single-finger representations, as measured in the average pattern dissimilarities, varied across regions (Fig. 2D). Previous imaging work (Martuzzi et al., 2014; Besle et al., 2014) has not accounted for this potential confound. Here we addressed this issue by computing the expected selectivity index for random tuning and for highly selective tuning, given the signal reliability in each region and each participant (see Methods). We then normalized the measured selectivity coefficients for each participant and region separately, with 0 indicating random tuning and 1 indicating highly selective tuning for a single finger only.

After carefully controlling for signal reliability across regions, we confirmed that voxels in BA 3b showed strong selectivity for single fingers (Fig. 3C), significantly more than expected from random tuning (one-sided t-test >0: t9=8.329, p=8.01e-6). In comparison, more posterior subregions of S1 were more broadly tuned, indicated by significantly lower selectivity indices compared to BA 3b (BA 1: t9=3.166, p=0.0057, BA 2: t9=4.292, p=0.0010). Indeed, in BA 2, the finger selectivity coefficients did not differ from what would be expected assuming random tuning curves (t9=–0.029, p=0.5112). Moving anterior relative to BA 3b, voxels were less selective in BA 3a (t9=3.900, p=0.0018).

Selectivity indices in M1 were significantly lower than in BA 3b, for both posterior (BA 4p, t9=6.944, p=3.366e-5) and anterior portions (BA 4a, t9=4.177, p=0.0012). Rathelot and Strick (2009) proposed that “new M1” (BA 4p) is more essential for individuated finger movements than “old M1” (BA 4a), from which one may predict that BA 4p should show more selective single-finger representation. To test this prediction, we contrasted BA 4p and BA 4a, which may approximate old and new M1, respectively. Contrary to this prediction we found no difference in the average selectivity coefficients between these regions (t9=−0.991, p=0.8262). Taken together, however, our analyses confirm the idea that sensory information from individual fingers is spatially quite segregated in BA 3b, but then continuously intermixes when moving more anterior or posterior in the sensorimotor cortex.

### Interactions between finger activity patterns explain spatial complexity of multifinger patterns

Having established that somatosensory inputs from different fingers heavily overlap in regions of the sensorimotor cortex, we then asked how somatosensory inputs are integrated across fingers. As a first step, we inspected the spatial activity patterns evoked during multi-finger stimulation. Figure 4A shows the activity patterns from two exemplary participants during stimulation of the index finger, the little finger, or during stimulation of both fingers. The spatial patterns evoked by stimulating each finger had a relatively focal activation point in BA 3b. For the combined stimulation, we can see two areas of activation, one corresponding to the region active for the index finger, and the other corresponding to the region active during for the little finger. This suggests that the representation of inputs from multiple fingers in BA 3b may be linear, simply reflecting the superposition of activity caused by the stimulation of the individual fingers. We would expect such linearity if the inputs from different fingers do not interact with each other.

**Figure 4.**
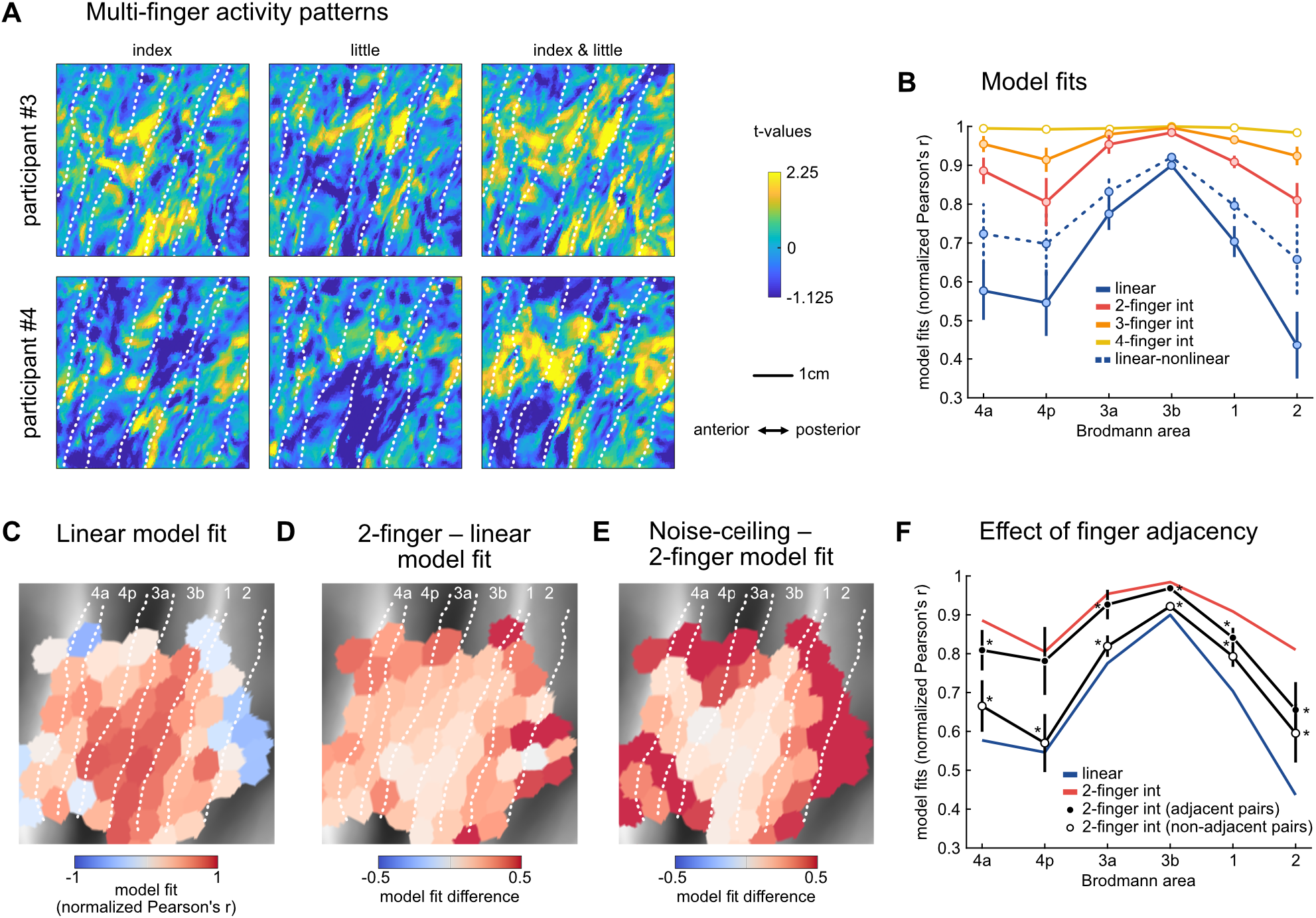
Multi-finger activity patterns in sensorimotor cortex. (**a**) Exemplar activity patterns from the two participants displayed in Figure 3A. (**b**) Representational model fits were normalized to the null model (0) and the noise-ceiling (1) in each region in each participant. Dots reflect the mean and error bars reflect SEM across participants per region. (**c**) Surface map of the linear model fit (median across participants) in tessels where there were significant differences between all finger combination patterns (average paired dissimilarities between finger combination patterns ≥0.005). (**d**) Difference between the fits of the 2-finger interaction model and the linear model in each tessel (median across participants). (**e**) Difference between the noise-ceiling and the fit of the 2-finger interaction model in each tessel. Note that the colour scale for C is different than that for D and E. (**f**) The full 2-finger interaction model (red line) was compared to a model that only contained the adjacent (black markers) or the non-adjacent (white markers) finger-pairs. Asterisks denote significantly lower model fits compared to using all two-finger interactions (two-sided paired t-test, *p*<0.05).

In contrast, the multi-finger spatial pattern in BA 1 and BA 2 appeared more complex, with clusters of activity emerging during simultaneous stimulation that were not present when either finger was stimulated individually. Given that the neural populations representing each finger appeared to be more overlapping in these same regions, the complexity of the spatial patterns suggests the presence of an interaction between fingers.

To test this idea formally, we fit a series of representational models to the activity patterns in each participant and region. These encoding-style models were fit to the activity patterns across all voxels in a region during single- and multi-finger stimulation, and then evaluated by their ability to predict multi-voxel activity patterns measured during an independent test run (see Methods). To meaningfully compare model fits across regions with different signal-to-noise levels, we normalized them to the performance of a null model and a noise-ceiling model. The null model predicted that the overall activity would increase when more fingers are stimulated, but that the activity patterns themselves would not differ between finger combinations. The noise-ceiling model was fit by estimating a unique pattern for each finger combination from the data (i.e., the model allowed for any arbitrary non-linearity). The model fits were then normalised between the null model (0) and noise-ceiling model (1), to express how much of the replicable finger-specific variation in the activity patterns each model could explain.

Based on the observations in BA 3b, we first examined to what degree multi-finger patterns were simply the sum of the constituent single-finger patterns. The predictive performance of this linear model was significantly better than that of the null model across the sensorimotor cortex (region x model ANOVA, main effect of model: *F_(1,9)_*=590.662, *p*=1.618e-9; see Fig. 4B), indicating that the linear model captured some reliable aspects of the spatial activity patterns. Furthermore, the normalized linear model fit varied across regions (region x model ANOVA, interaction effect: *F_(5.45)_*=7.308, *p*=4.385e-5). The best fit was observed in BA 3b, with significantly lower fits in all other regions (all *t_9_*≥4.139, all *p*≤0.0025, evaluated at a Bonferroni corrected *α*-value=0.01) except BA 3a (*t_9_*=2.822, *p*=0.0200), where the difference was not significant after correction. Importantly, in all regions the linear model predicted the data significantly worse than the noise-ceiling model (all *t_9_*≥5.318, all *p*≤0.0005), indicating that there were systematic non-linearities in the multi-finger activity patterns that could not be explained by the linear model.

To visualize more generally how the linear model fit across the sensorimotor cortex in a region-blind manner, we applied the same model fitting to data from regularly tessellated regions (Fig. 4C, see Methods). This yielded similar results, with good fits in BA 3b and increasing non-linearities in regions anterior and posterior to it (denoted by the drop in performance of the linear model).

We then considered the possibility that non-linearities in how the activity patterns for single fingers combine would arise from the interaction of pairs of fingers, perhaps via local surround-inhibition or divisive normalization between two finger representations. Therefore, we created a two-finger interaction model, which explained all patterns as the sum of the component single-finger patterns, as well as their two-finger interactions (see Methods). Across all regions, this two-finger interaction model predicted left-out data significantly better than the linear model (region × model ANOVA, main effect of model: *F_(1,9)_*=209.851, *p*=1.526e-7).

The amount of variance explained by these two-finger interactions, however, differed across regions (Fig. 4D). While the two-finger interactions lead to a small gain in predictive performance in BA 3b (8.44±0.83%), the gain was over four times larger in BA 2 (37.37±4.41%). Indeed, the region x model interaction effect was highly significant (*F_(5,45)_*=9.753, *p*=2.320e-6). This indicates that a larger proportion of the pattern variance could be explained in regions outside of BA 3b when including interaction effects between pairs of fingers.

### Interactions do not only arise between adjacent fingers

Do the non-linear interactions between fingers described above arise mostly between adjacent fingers or do interactions also arise between spatially distant fingers of the hand? Previous work has shown that stimulating adjacent fingers leads to lower activity compared to non-adjacent fingers, which has been interpreted as evidence that adjacent fingers interact more than non-adjacent fingers (Biermann et al., 1998; Friedman et al., 2008; Hsieh et al., 1995; Ishibashi et al., 2000; Lipton et al., 2010). However, whether adjacent finger interactions are stronger across all regions of the sensorimotor cortex is not known. We investigated this by fitting two variants of the full two-finger interaction model, either including only the interaction terms for either adjacent or non-adjacent finger-pairs. Using only the non-adjacent pairs resulted in significantly lower model performance in all regions compared to using all finger-pairs (all *t_9_*≤−5.609, all *p*≤0.0003; see Fig. 4F). When we used the adjacent finger pairs, the model performance was not significantly reduced in BA 4p (*t_9_*=–0.605, *p*=0.5600), BA 4a (*t_9_*=−2.793, *p*=0.0210), and BA 3a (*t_9_*=−1.318, *p*=0.2199) when correcting for multiple comparisons (*α*-value=0.0083). In contrast, the fits in BA 3b, BA 1, and BA 2 were significantly lower (all *t_9_*≤−4.611, all *p*≤0.0013). Taken together, this suggests that in posterior regions of the sensorimotor cortex, interactions between both adjacent and non-adjacent finger-pairs were important in explaining the multi-finger activity patterns. Furthermore, a significant region x model interaction (*F_(5,45)_*=3.199, *p*=0.0148) indicated that the effect of finger adjacency differed across regions. In BA 2, the predictive power of adjacent or non-adjacent interactions was comparable (twosided paired t-test: *t_9_*=–1.403, *p*=0.1941), whereas non-adjacent interactions were significantly less important in all other subregions (all *t_9_*≤−3.090, all *p*≤0.0130). This suggests that BA 2 shows strong interactions between finger-pairs irrespective of finger adjacency.

### Complexity of finger interactions increases along sensorimotor cortex

Thus far, we have demonstrated that population activity across sensorimotor cortex strongly represents two-finger interactions. However, in order to provide the neural substrate necessary to skillfully manipulate an object held in the entire hand, the sensorimotor system needs to be able to detect specific patterns of stimulation across all fingers. Therefore, we should find evidence for integration of information across more than two fingers.

In BA 3b and BA 3a, the two-finger interaction model provided a good fit to the multi-finger activity patterns. In these regions, the model performance of the two-finger interaction model was very close to the noise-ceiling model, accounting for 98.42±0.38% and 95.37±2.40% of the reliable pattern variance, respectively. While a small significant difference remained in BA 3b (two-sided paired t-test: *t_9_*=4.162, *p*=0.0024), the two-finger interaction model explained the activity patterns as well as the noise-ceiling model in BA 3a (*t_9_*=1.916, *p*=0.0877). Thus, neural populations in BA 3b and BA 3a do not appear to provide a unique, and hence linearly separable, representation of all possible multi-finger combinations.

In contrast, the predictive performance of the two-finger interaction model was still lower than the noise-ceiling in the other regions (all *t_9_*≥3.142, all *p*≤0.0119; see Fig. 4E). We therefore considered the interactions of three fingers in our models (see Methods). By including three-finger interactions, we were able to explain the activity patterns as well as the noise-ceiling model in BA 4a (*t_9_*=2.183, *p*=0.0569). In BA 4p, BA 1, and BA 2, however, performance was still significantly lower than the noise-ceiling (all *t_9_*≥2.731, *p*≤0.0232). Only after including four-finger interactions were we able to fully explain the activity patterns in these remaining regions (all *t_9_*≤2.154, *p*≥0.0597). This suggests that the interactions in the most anterior and posterior regions of the sensorimotor cortex are more complex, involving non-linear interactions between three or more fingers. Therefore, our results appear to indicate that BA 1, BA 2, and also BA 4 integrate sensory information arriving from multiple fingers to create a unique representation of specific patterns of multi-finger stimulation.

### Finger interactions do not reflect a general suppression of activity

There is, however, an alternative and relatively simple mechanism that could give rise to the poor performance of the linear model in subregions of the sensorimotor cortex. Specifically, it may be the case that the single-finger activity patterns combine linearly, but that the overall activity in each region is scaled in a non-linear fashion. Such non-linear scaling could arise from divisive normalization of the overall activity within the region, or from non-linearities between neural activity and the BOLD signal.

To test this, we expanded the linear model to allow for non-linear scaling of overall activity (see Methods). Indeed, this linear-nonlinear model provided a significantly better fit than the original linear model in all BA regions (two-sided paired t-test: all *t_9_*≥6.105, all *p*≤0.0002; see Fig. 4B). This alone should not be too surprising, given that the average activity did not scale linearly with the number of fingers stimulated (Fig. 2B). Importantly, however, the predictive performance of the two-finger interaction model remained significantly better than that of the linear-nonlinear model in BA 3a, BA 3b, BA 1, and BA 4 (all *t_9_*≥3.395, all *p*≤0.0079). Although this difference was not significant in BA 2 after applying Bonferroni correction for multiple comparisons (*t_9_*=3.291, *p*=0.0094, *α*-value=0.0083), the two-finger interaction model still accounted for 15.23α4.63% more pattern variance in this region. Furthermore, compared to the higher-order interaction models, the linear-nonlinear model performed substantially worse in BA 2 (vs. three-finger: *t_9_*=−2.837, *p*=0.0195; vs. four-finger: *t_9_*=−3.715, *p*=0.0048), and more generally across the sensorimotor cortex (Fig. 4B). Therefore, the non-linearities captured by our multi-finger interaction models likely reflect complex interactions that arise between specific sets of finger patterns, rather than simply reflecting a general non-linear scaling of activity across the region.

## Discussion

In this study, we investigated how somatosensory information coming from the fingers is integrated in different areas of the sensorimotor cortex. We hypothesized that to guide skillful object manipulation, the sensorimotor system needs to be able to detect relatively arbitrary combinations of sensory inputs across fingers, requiring non-linear integration of any pair, triplet, and quadruplet of fingers. We reported that voxels in BA 3b tend to be selectively tuned to the inputs from a single finger, whereas regions anterior and posterior show less finger-specificity, even after we controlled for differences in signal to noise. In previous work, this broader tuning to multiple fingers has often been interpreted as evidence for finger integration (Iwamura et al., 1993; Martuzzi et al., 2014). However, spatial overlap itself only suggests that individual fingers are represented in overlapping neural populations– it does not necessarily mean that information from different fingers is integrated. By using the full set of multi-finger combinations and representational model analyses, we could show that the multi-finger patterns could not be explained by a simple linear combination of the single-finger patterns. Rather, most regions showed clear non-linearities, which not only reflected interactions between pairs of finger-pairs, but by any combination of multiple fingers.

An important strength of our experimental design was that it allowed us to test whether the observed non-linearities really reflected integration of information across the fingers. Alternatively, a relatively simple explanation for our results is that the activity patterns caused by single finger stimulations are simply summed but that the overall activity is then suppressed in a non-linear fashion. Previous studies have been unable to distinguish between these two explanations, as they used very few multi-finger combinations, making it difficult to dissociate global non-linear activity suppression from unique non-linear interactions (Gandevia et al., 1983; Hsieh et al., 1995; Biermann et al., 1998; Ishibashi et al., 2000; Hoechstetter et al., 2001; Ruben et al., 2006; Lipton et al., 2010; Brouwer et al., 2015). Having stimulated all possible multi-finger combinations, we had sufficient leverage to distinguish between these two possibilities and were able to rule out simple global suppression. That is, the interactions we report in this paper likely reflect rich, non-linear integration of sensory inputs from the fingers.

The fit of the linear combination model was greatest in BA 3b, where neurons have receptive fields that are largely restricted to single fingers (Sur, 1980; Iwamura et al., 1993) and preferentially code for tactile features that can be extracted from local spatial regions such as stimulus edge orientation (Hyvärinen & Poranen, 1978a; Bensmaia et al., 2008). However, this is not to say that multi-finger integration was entirely absent in BA 3b. Indeed, consistent with previous work (Reed et al., 2008, 2010; Lipton et al., 2010; Thakur et al., 2012), we found significant finger-pair integration in BA 3b. Interactions were stronger between adjacent fingers, indicating that the majority of integration that occurs in BA 3b is across spatially close distances, as previously reported (Reed et al., 2008).

Moving posterior from BA 3b to BA 2, we observed progressively more complex multi-finger interactions, with evidence for non-linear interactions of all possible multi-finger combinations in BA 2. This complexity matches the changes in tactile feature preference of individual neurons, shifting from local tactile features like edge orientation (Bensmaia et al., 2008) to higher-order features that span multiple fingers like object curvature (Yau, Connor, & Hsiao, 2013). The interactions also occurred for finger pairs of increasing spatial distance. Indeed, the interactions between adjacent and non-adjacent fingers were equally strong in BA 2. Such broad spatial integration is important for extracting spatially invariant higher-order tactile features of an object (Yau et al., 2016). Together, these observations provide empirical support for the hypothesis that tactile inputs from the hand are progressively elaborated along the sensory cortical pathway (Hyvärinen & Poranen, 1978b; Phillips, Johnson, & Hsiao, 1988; Iwamura, 1998).

We also examined sensory processing in the hand area of M1, which has commonly been ignored in previous work. Although BA 4 is traditionally viewed as a motor area, it receives substantial inputs from the somatosensory thalamus (Jones, 1975; Darian-Smith & Darian-Smith, 1993) and from various areas of S1 (Ghosh, Brinkman, & Porter, 1987). Therefore, neural populations in this region may also be involved in integrating tactile inputs from the fingers, perhaps for rapid behavioural responses to object displacements (Crevecoeur et al., 2017; Hernandez-Castillo et al., 2020). Our results demonstrate that there were finger interactions in BA 4, and the strength of these interactions were comparable to those in BA 2. However, it is not clear whether these interactions arose specifically within BA 4 or reflect inputs from BA1 or BA 2

What benefit does non-linear integration of information across fingers provide? From an ethological standpoint, non-linear finger integration allows for a more flexible mapping between sensory inputs and motor responses. For example, consider the scenario where you are holding a cup in your hand. Any movement of the cup across your fingers, be it slipping downward out of your hand or rotating outward out of your hand, needs to be countered with an increase in grip force if the goal is to stabilize the cup in your grasp (Cole & Abbs, 1988). However, the appropriate response at more proximal muscles will be quite different in these circumstances. Downward movement of the cup will largely require activation of muscles that radially deviate the wrist and flex the elbow whereas outward cup rotation will largely require activation of muscles that deviate the wrist but not flex the elbow. In general, the appropriate muscle recruitment pattern cannot be determined by a linear readout of inputs from each individual finger, since this could produce unnecessary or even counterproductive responses. Only non-linear integration of the slip signals would allow the mapping of these different patterns of sensory inputs to the appropriate patterns of muscle recruitment.

Our results are in agreement with recent evidence from multi-whisker stimulation studies in rodents, where specific combinations across whiskers are uniquely represented by neural populations in the rodent barrel cortex (Laboy-Juárez et al., 2019; Lyall et al., 2021). Like the human hand, the rodent whisker system has evolved to support complex spatial-temporal interactions with the environment. Together with our current results, this suggests that non-linear integration between somatosensory inputs occurs when the detection of complex sensory patterns is ethologically significant.

In our experiment, we required participants to remain attentive to the finger stimulation. Processes of selective attention have been shown to modulate neural firing rates in response to finger stimulation (Hsiao, O’Shaughnessy, & Johnson, 1993). To what degree are our findings caused by raw sensory input, and to what degree did our specific mismatch detection task influence somatosensory processing? Although our task required the comparison of the pattern of stimulation across fingers to a visual stimulus, the visual stimulus was only presented after the somatosensory stimulation. At the moment of finger stimulation, participants had no expectation as to which finger combination would be stimulated. Therefore, the initial and dominant response observed in the fMRI data should reflect bottom-up somatosensory processing. More importantly, the mismatch task did not require integration of sensory information across fingers. Accurate performance could be achieved by simply judging sensory information from each finger in an independent manner. Therefore, if our mismatch task did induce any bias in the observed finger representations, the bias is more likely to be towards an independent representation of finger-specific inputs.

In general, it is very likely that task demands will influence how sensory information from the hand is processed (to some degree). Indeed, neural populations in S1 are modulated by inputs from M1 (Goldring et al., 2014), and the neural state of S1 is strongly influenced by the planning of upcoming actions (Ariani, Pruszynski, & Diedrichsen, 2022; Gale, Flanagan, & Gallivan, 2021). Such modulation is important, as the processing requirements of somatosensory information depends on the task at hand. For example, the reaction to object slip depends not only on the direction of the slipping object (Häger-Ross, Cole, & Johansson, 1996), but also on the perceived physical properties of the object (i.e., how “object-like” the simulation is, Ohki, Edin, & Johansson, 2002) and the behavioural goal (Hernandez-Castillo et al., 2020). We may therefore expect that, in order to provide support for flexible sensory-motor mapping, the way that information is integrated across fingers changes with behavioral context. Thus, the next challenge is to probe how such top-down influences alter the integration of somatosensory inputs across fingers.

## Acknowledgements

The work was supported by a Discovery Grant from the Natural Sciences and Engineering Research Council (NSERC, RGPIN-2016-04890) to JD. Functional imaging costs were partly supported by a Platform Support Grant from Brain Canada and the Canada First Excellence Research fund (CFREF, BrainsCAN). SA was supported by a doctoral scholarship from NSERC (PGSD3-519263-2018). JP is supported by the Canada Research Chairs program.

